# Role of posterodorsal medial amygdala urocortin-3 in pubertal timing in female mice

**DOI:** 10.1101/2022.01.27.477996

**Authors:** Deyana Ivanova, Xiao-Feng Li, Yali Liu, Caitlin McIntyre, Cathy Fernandes, Geffen Lass, Lingsi Kong, Kevin T O’Byrne

## Abstract

Post-traumatic stress disorder impedes pubertal development and disrupts pulsatile LH secretion in humans and rodents. The posterodorsal sub-nucleus of the medial amygdala (MePD) is an upstream modulator of the hypothalamic gonadotropin-releasing hormone (GnRH) pulse generator, pubertal timing, as well as emotional processing and anxiety. Psychosocial stress exposure alters neuronal activity within the MePD increasing the expression of Urocortin3 (Ucn3) and its receptor corticotropin-releasing factor type-2 receptor (CRFR2) while enhancing the inhibitory output from the MePD to key hypothalamic reproductive centres. We test the hypothesis that psychosocial stress, processed by the MePD, is relayed to the hypothalamic GnRH pulse generator to delay puberty in female mice. We exposed C57Bl6/J female mice to the predator odor, 2,4,5-Trimethylthiazole (TMT), during pubertal transition and examined the effect on pubertal timing, pre-pubertal LH pulses and anxiety-like behaviour. Subsequently, we virally infected Ucn3-cre-tdTomato female mice with stimulatory DREADDs targeting MePD Ucn3 neurons and determined the effect on pubertal timing and pre-pubertal LH pulse frequency. Exposure to TMT during pubertal development delayed puberty, suppressed pre-pubertal LH pulsatility and enhanced anxiety-like behaviour, while activation of MePD Ucn3 neurons reduced LH pulse frequency and delayed puberty. Early psychosocial stress exposure decreases GnRH pulse generator frequency delaying puberty while inducing anxiety-behaviour in female mice, an effect potentially involving Ucn3 neurons in the MePD.

## Introduction

In humans post-traumatic stress disorder (PTSD) is associated with altered pubertal timing and the development of anxiety disorders (1,2). Predator odor exposure is a classic model for PTSD in rodents (3) and has been shown to delay puberty (4), suppress luteinizing hormone (LH) pulse frequency (5) and inhibit the pre-ovulatory LH surge (6). The gonadotropin-releasing hormone (GnRH) pulse generator that controls the hypothalamus-pituitary-gonadal (HPG) axis and pubertal development is restrained in the juvenile, but reactivated at a critical time point with increased frequency leading to the initiation of puberty (7). However, the neural mechanisms controlling pubertal timing are not fully established.

Kisspeptin (kiss1) is known to be a major gatekeeper of pubertal onset. An absence of the *kiss1* gene and kiss1 receptor (kiss1r) in mice and humans leads to a loss of gonadal maturation and a lack of pubertal onset (8,9), and treatment with kisspeptin can reactivate the HPG axis during the juvenile hiatus in monkeys and rats (10,11). Kiss1 neurons are located in the anteroventral periventricular nucleus (AVPV) and arcuate nucleus (ARC) of the hypothalamus in mice (12). The ARC kiss1 neurons, known as KNDy because they co-express neurokinin B (NKB) and dynorphin A (Dyn), are a critical component of the GnRH pulse generator (12–14). The activation of KNDy neurons induces pulsatile release of kiss1 acting on GnRH dendrons in the median eminence to stimulate pulsatile GnRH secretion (15). The AVPV kiss1 neurons project to GnRH cell bodies and proximal dendrites and are known to control the preovulatory LH surge and ovulation in rodents (16). In the juvenile, a neurobiological break exerted on the ARC GnRH pulse generator suppresses pulsatile kiss1 release in the median eminence, however the nature of this upstream inhibition of the GnRH pulse generator, which is released at the end of the juvenile phase to trigger puberty is unknown (7).

The amygdala, a stress-sensitive part of the limbic brain, is involved in processing anxiety and fear as well as modulating pubertal timing and the HPG axis. Lesioning of the amygdala advances menarche in female rhesus macaques (17), whereas electrical stimulation of this region delays puberty in rats (18). Specific lesioning of the posterodorsal sub-nucleus of the medial amygdala (MePD) advances puberty in rats (19), suggesting this region may play a key role in exerting an inhibitory break on pubertal timing. The MePD sends GABAergic projections to various hypothalamic reproductive centres (20,21) and has been shown to project directly to ARC KNDy neurons, although their neurochemical phenotype is undetermined (22,23). Interestingly, the MePD contains a kiss1 neuronal population (24) and antagonism of MePD kiss1 delays puberty, disrupts estrous cyclicity and the LH surge in rats (25). Moreover, selective optogenetic stimulation of MePD kiss1 neurons increases LH pulse frequency in mice (26). The MePD is highly responsive to psychosocial stress, exhibiting a distinct firing pattern in response to predator odor (27) and recently cat odor was found to delay puberty, disrupt estrous cyclicity and induce a fear response in rats (4). The MePD may be a central hub involved in the integration of external olfactory and anxiogenic signals with the GnRH pulse generator.

Central administration of the stress neuropeptide corticotrophin releasing factor (CRF) dose-dependently delays puberty, while CRF-receptor antagonism advances puberty in rats (28). Urocortin-3 (Ucn3), a member of the CRF family and an endogenous ligand for CRF type 2 receptors (CRFR2) is found in abundance in the MePD (29). Psychosocial stressors, including social defeat and restraint, increase cfos expression in CRFR2 positive neurons and increase Ucn3 mRNA expression in the MePD of rodents (30). Moreover, we have recently shown that MePD Ucn3 mediates predator odor and restraint stress-induced suppression of pulsatile LH secretion in mice (5).

In this study, we aimed to determine whether chronic predator odor stress exposure suppresses pre-pubertal LH pulse frequency and delays pubertal timing in female mice. Additionally, we assessed social, anxiety and fear-like behaviour as measures of psychosocial stress in predator odor-exposed pre-pubertal female mice. Finally, we aimed to determine whether MePD Ucn3 signalling is involved in modulating pre-pubertal LH pulse frequency and pubertal timing in female mice.

## Materials and Methods

### Mice

Breeding pairs of C57Bl6/J mice were purchased from Charles River Laboratories International, Inc. For Ucn3-cre mice, cryopreserved sperm of strain (Tg(Ucn3-cre)KF43Gsat/Mmucd; congenic on C57BL/6 background) was acquired from MMRRC GENSAT. Breeding pairs of heterozygous transgenic Ucn3-cre mice were recovered by insemination of female C57Bl6/J mice at King’s College London. Genotyping of Ucn3-cre mice was performed using PCR to detect heterozygosity. Heterozygous Ucn3-cre mice were bred with cre-activated tdTomato reporter mice (strain B6.Cg-Gt(ROSA)26Sortm9(CAG-tdTomato)Hze/J; congenic on C57BL/6 background obtained from The Jackson Laboratory, Bar Harbor, ME, USA) to obtain Ucn3-cre-tdTomato mice, as previously described (5,29). Litter size was reduced to 6-8 pups 2-3 days after birth (on pnd 2-3) to standardise body weight, which can alter pubertal development. Mice aged between 21 to 45 days were group housed in individually ventilated cages equipped with wood-chip bedding and nesting material with food and water ad libitum and sealed with a HEPA-filter at 25 ± 1 °C in a 12:12 h light/dark cycle, lights on at 07:00 h. All procedures were carried out following the United Kingdom Home Office Regulations and approved by the Animal Welfare and Ethical Review Body Committee at King’s College London.

### Stereotaxic adeno-associated-virus injection

All surgical procedures were carried out with aseptic conditions and under general anaesthesia using ketamine (Vetalar, 100 mg/kg, i.p.; Pfizer, Sandwich, UK) and xylazine (Rompun, 10 mg/kg, i.p.; Bayer, Leverkusen, Germany). The mouse brain atlas of Paxinos and Franklin (31) was used to obtain target coordinates for the MePD (2.15 mm lateral, −1.25 mm from bregma, at a depth of −5.30 mm below the skull surface) of pre-pubertal mice. Mice at pnd 14 were temporarily taken from their mother and secured in a David Kopf stereotaxic frame (Kopf Instruments), a small skin incision was made to reveal the skull and two small holes were drilled above the location of the MePD. Bilateral stereotaxic viral injections of the stimulatory adeno-associated-virus (AAV) carrying the DIO-hM3D-mCitrine, DREADD, construct (AAV-hSyn-DIO-HA-hM3D(Gq)-IRES-mCitrine, 3×10^11^ GC/ml, Serotype:5; Addgene) was administered intra-MePD using the robot stereotaxic system (Neurostar, Tubingen, Germany), performed for the targeted expression of DIO-hM3D-mCitrine in MePD Ucn3 neurons in Ucn3-cre-tdTomato mice. AAV-hSyn-DIO-HA-hM3D(Gq)-IRES-mCitrine (150 nl) was bilaterally injected into the MePD using a 2-μl Hamilton micro syringe (Esslab, Essex, UK) over 10 min and the needle was left in position for a further 5 min then slowly lifted over 2 min. Mice that were cre-positive received the AAV-hM3D injection (test mice) or a control virus AAV-YFP (Addgene) (control mice) where the control virus does not contain the DIO-hM3D-mCitrine construct. The mice were placed back with the mother for a further 7 days until wean day (pnd 21). Nine Ucn3-cre-tdTomato mice received the AAV-hM3D injection, 6 Ucn3-cre-tdTomato mice received the control AAV-YFP, 4 Ucn3-cre-negative mice without surgery received only CNO and 3 Ucn3-cre-tdTomato mice without surgery did not receive CNO.

### Chronic pre-pubertal predator-odor exposure and puberty evaluation

Female pups were weaned on pnd 21 and separated into control and test groups. To investigate the effects of chronic pre-pubertal psychosocial stress mice were removed from their home cages, singly housed in new cages and left to habituate for 10 min. After the habituation period, the mice were exposed to 12 μl of 2,4,5-Trimethylthiazole (TMT; synthetic extract of fox urine; ≥98% purity; Sigma-Aldrich, UK) pipetted on a small circular piece of filter paper in a petri dish placed in the centre of the cage for 20 min. The mice were exposed to TMT daily, at random time points, for 14 days (from pnd 21 to 35). Control mice were exposed to filter paper soaked with 12 μl of ddH2O water. To rule out the possibility that physiological and behavioural responses in TMT-exposed mice resulted from novelty of scent we repeated the experiment with 85 mM of ethyl vanillin (≥98% purity; Sigma-Aldrich UK) as a control scent in a separate cohort of mice. Mice were monitored from pnd 24 for vaginal opening (VO) and first estrous (FE) indicated by epithelial cell cornification. Once the occurrence of VO was detected, vaginal smears were taken to determine the exact day of FE.

### Pre-pubertal blood sampling

The effect of TMT exposure on pre-pubertal LH pulses was measured, on pnd 26 and 29. Mice were handled twice daily for 10 min to habituate to tail-tip blood collection for at least 7 days prior to blood sampling, as described previously (5). For LH measurement, 3 μl of blood was collected every 5 min for 80 min. Blood sampling was performed on pnd 26 and 29 during the period of daily TMT exposure (pnd 21-35). Blood samples were collected at least 2 h before TMT-exposure on these days. Blood collection was performed between 09:00-12:00 h. For Ucn3 DREADDs experiments, blood sampling was performed on pnd 29 during the period of daily CNO administration (pnd 21-35), as described above. Blood collection was performed between 09:00-12:00 h.

### Behavioural tests

#### Light Dark Box

The same cohort of C57Bl6/J female mice were transported to a new room (lights off) and left to habituate for 30 min. Mice were placed in the centre of the light compartment facing towards the dark compartment. The time spent in each compartment (entry with all 4 limbs) was recorded manually over a period of 5 min by two independent observers. The LDB test was carried out on pnd 19, 27 and 40. Behaviour testing performed during the period of daily TMT exposure (pnd 21-35) and was measured before the exposure to TMT on that day. This applies to all behaviour tests performed during the TMT exposure period. All behaviour tests were performed between 11:00-13:00 h.

#### Social interaction with familiar conspecific

The same cohort of C57Bl6/J female mice were transported to a new dimly lit room and left to habituate for 30 min. Experimental mice and a same sex familiar conspecific (similar age, weight and strain) were placed simultaneously into the test arena (a new cage with clean wood-chip bedding). The time spent following, sniffing, grooming and mounting the conspecific was monitored and recorded manually over a period of 5 min by two independent observers. The social interaction test was carried out on pnd 19, 27 and 40.

#### Elevated Plus maze

The same cohort of C57Bl6/J female mice were transported to a new room (lights on) and left to habituate for 30 min. Mice were placed in the centre of the maze facing towards the closed arms. The time spent in each part of the maze (open, centre and closed; entry with all 4 limbs) was recorded manually over a period of 5 min by two independent observers. The EPM test was carried out on pnd 20, 28 and 41.

### Chronic DREADD activation of Ucn3 neurons in the MePD

For DREADDs experiments, stock solution of Clozapine-N-oxide (CNO) (Tocris Bio-techne, Abingdon, UK) was made by dissolving 5 mg of CNO in 1 ml of 0.9% sterile saline solution and stored at 4 °C. CNO was made fresh daily and administered via drinking water at a final concentration of 0.5 mg CNO/ kg, as described previously (32) for 14 days from pnd 21-35. Mice were monitored from pnd 22 for vaginal opening (VO) and first estrous (FE), as described above.

### Validation of AAV injection

On pnd 50, Ucn3-cre-tdTomato mice were anaesthetised with a lethal dose of ketamine. Transcardial perfusion was performed with heparinised saline for 5 min followed by ice-cold 4% paraformaldehyde (PFA) in phosphate buffer (pH 7.4) for 15 min with a pump (Minipuls, Gilson, Villiers Le Bel, France). Brains were immediately collected and fixed in 15% sucrose in 4% PFA at 4 °C and left to sink. Brains were then transferred to 30% sucrose in phosphate-buffered saline (PBS) and left to sink. Brains were snap-frozen in isopropanol on dry ice and stored in −80ºC. Every third coronal brain section, 30-μm/section, was collected using a cryostat (Bright Instrument Co., Luton, UK) through-out the MePD region corresponding to −1.34 mm to −2.70 mm from bregma. Brain sections were mounted on microscope slides, air dried and covered with ProLong Antifade mounting medium (Molecular Probes, Inc. OR, USA). The number of tdTomato labelled and AAV-hM3D-mCitrine infected Ucn3 neurons per side per slice in the MePD was quantified from 4 slices. For AAV-hM3D-mCitrine injected Ucn3-cre-tdTomato mice we determined whether Ucn3 neurons were infected in the MePD region by merging td-Tomato fluorescence of Ucn3 neurons with mCitrine fluorescence in the MePD. Images were taken using Axioskop 2 Plus microscope (Carl Zeiss) equipped with axiovision, version 4.7 (Carl Zeiss). Only data from animals with correct AAV injection were analysed.

### LH pulse detection and analysis

Blood samples were processed with a LH ELISA, as previously reported (33). Capture antibody (monoclonal antibody, anti-bovine LHβ subunit, AB_2665514) was purchased from Department of Animal Science at the University of California, Davis. Mouse LH standard (AFP-5306A) and primary antibody (polyclonal antibody, rabbit LH antiserum, AB_2665533) were obtained from Harbour-UCLA (California, USA). Secondary antibody (Horseradish-Peroxidase (HRP)-linked donkey anti-rabbit IgG polyclonal antibody, AB_772206) was purchased from VWR International (Leicestershire, UK). Inter-assay and intra-assay variations were 4.6% and 10.2%, respectively and the assay sensitivity was 0.0015 ng/mL. ODs of the standards were plotted against the log of the standard concentrations, non-linear regression to fit the points and parameters were extracted to calculate the concentration of LH (ng/ml) in blood samples, as previously described (33). The LH concentration at every time point of blood collection was plotted as a line and scatter graph using Igor Pro 7, Wavemetrics, Lake Oswego, OR, USA. DynPeak algorithm was used for the detection of LH pulses (34).

### Statistics

For TMT-exposed and DREADD injected female pre-pubertal mice, data obtained for VO and FE was analysed using RM one-way ANOVA. Data obtained for performance on LDB, SI and EPM, LH pulse frequency and body weight measure were analysed using RM two-way ANOVA. Statistics were performed using Igor Pro 7, Wavemetrics, Lake Oswego, OR, USA. Data was represented as mean ± SEM and +p<0.05, ++p<0.001 and +++p<0.0001 were considered to be significant.

## Results

### Chronic TMT exposure delays puberty onset

TMT exposure during the pubertal transition period, pnd 21 to 35, delayed FE without altering VO compared water and ethyl vanillin controls in C57Bl6/J female mice (Fig. 1, A and B; Control vs TMT, +++p<0.0001; TMT, n=14, water control, n=6, ethyl vanillin control, n=4). Data for water and ethyl vanillin exposed control mice were combined as control since there was no significant difference between the two control groups. Body weight was unaffected between the experimental groups (Fig. 1, C; Control vs TMT). The data from this study shows that predator-odor stress exposure during the pubertal transition delays pubertal onset.

**Figure 1.**
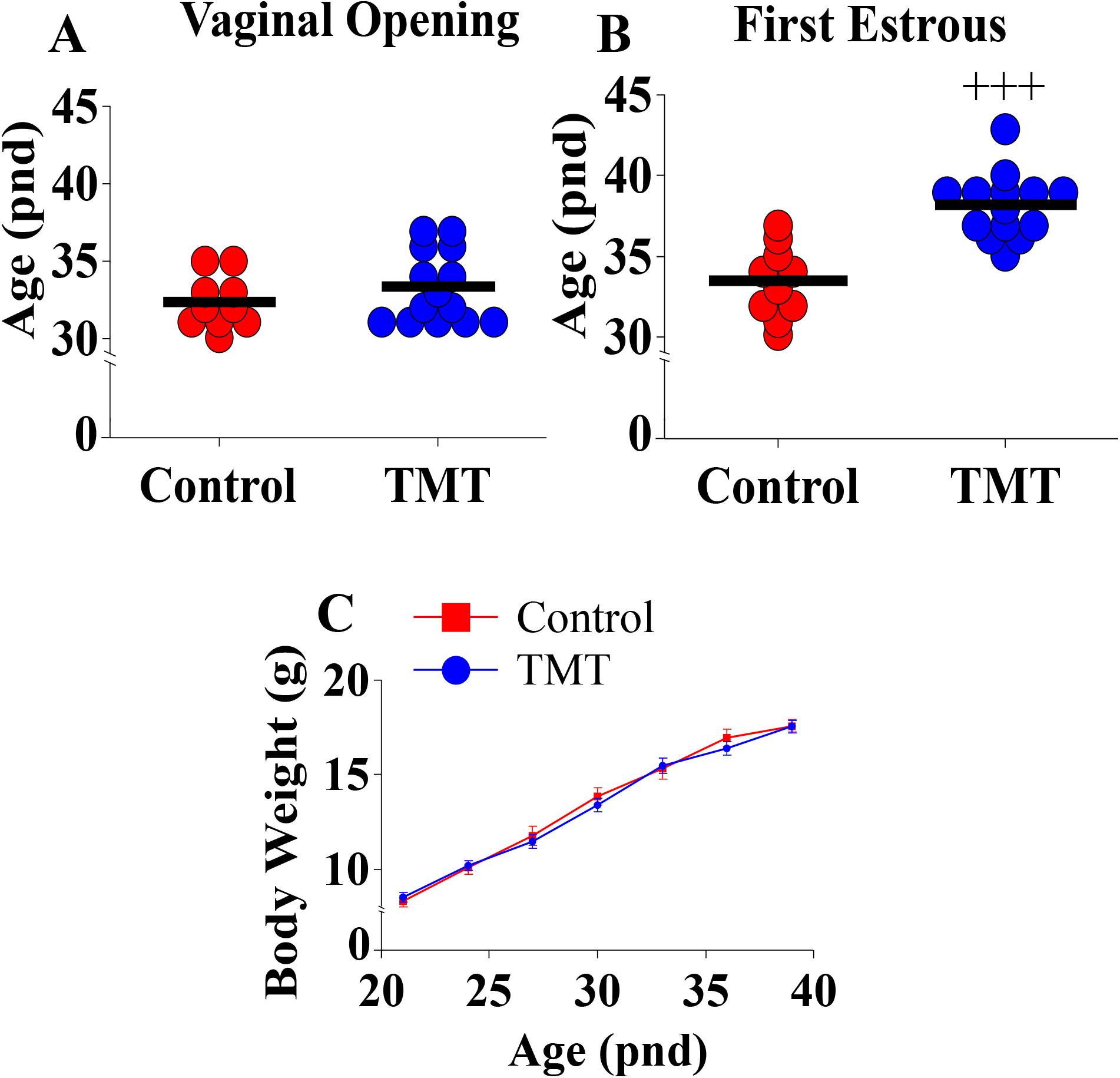
Female C57Bl6/J mice chronically exposed to 2,4,5-Trimethylthiazole (TMT) from post-natal day (pnd) 21 for 14 days showed delayed first estrous (FE) without affecting vaginal opening (VO) and body weight (BW). Effect of chronic TMT exposure on day of VO (A), FE and (C) BW weight gain between pnd 21 and 45. +++p<0.0001 Control group (combined water control: n=6; ethyl vanillin control: n=4) vs TMT group (n=14).

### Chronic TMT exposure decreases pre-pubertal LH pulse frequency

TMT exposure during the pubertal transition period, pnd 21 to 35, suppressed pre-pubertal LH pulse frequency on pnd 26 and 29 compared to water and ethyl vanillin controls in C57Bl6/J female mice (Fig. 2, A-E; Control vs TMT, ++p<0.001; TMT, n=8, water control, n=6, ethyl vanillin control, n=3). On pnd 29, the average LH pulse frequency for the control group tended to increase compared to the control group on pnd 26 potentially marking an acceleration of GnRH pulse generator activity approaching the onset of puberty (Fig. 2, A, C and E). The results of this experiment are summarised in the figure 2E. Data for water and ethyl vanillin exposed control mice were combined as control since there was no significant difference between the two control groups. The data from this study shows that predator-odor stress exposure during the pubertal transition inhibits pre-pubertal LH pulsatility.

**Figure 2.**
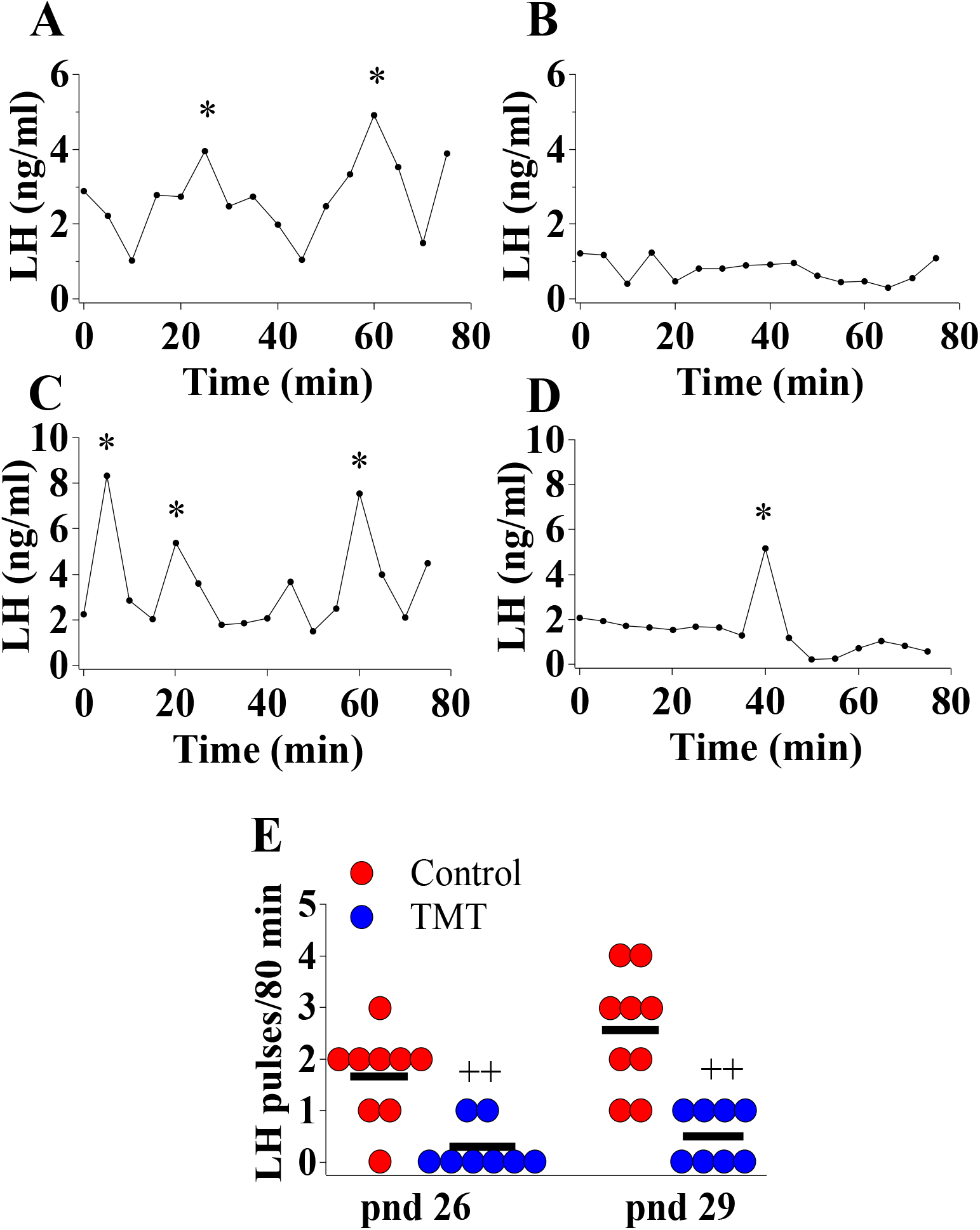
Female C57Bl6/J mice chronically exposed to 2,4,5-Trimethylthiazole (TMT) from post-natal day (pnd) 21 for 14 days showed suppressed pre-pubertal pulsatile luteinising hormone (LH) secretion on pnd 26 and 29. Representative LH pulse profile with (A) control mouse on pnd 26, (B) TMT-exposed mouse on pnd 26, (C) control mouse on pnd 29 and (D) TMT-exposed mouse on pnd 29. (E), Summary of LH pulse frequency on pnd 26 and 29 of control and TMT-exposed group during the 80 min blood sampling period. LH pulses detected by the DynePeak algorithm are indicated with an asterisk located above each pulse on the representative LH pulse profiles. ++p<0.001 control group (combined water control: n=6; ethyl vanillin control: n=3) vs. TMT-exposed group (n=8) on pnd 26 and 29.

### Chronic TMT exposure induced long-lasting anxiety and fear-like behaviour

We investigated anxiety and fear-like behaviour induced by TMT-exposure during the pubertal transition period, pnd 21 to 35 in in C57Bl6/J female mice. Anxiety and fear-behaviour were assessed using standard anxiety and fear tests, the LDB, SI and EPM, on pnd 19 or 20 (prior to day of weaning and TMT-exposure), pnd 27 or 28 (during the TMT exposure period) and pnd 40 or 41 (a week after termination of TMT-exposure). The first LDB, SI and EPM trial on pnd 19 or 20, prior to weaning, confirms no group difference in anxiety before the beginning of TMT-exposure (Fig. 3, A-C). TMT-exposed mice spent significantly less time in the light compartment of the LDB as well as socialising with a familiar conspecific compared to the control group on pnd 27, but not on pnd 40 (Fig. 3, A, B; Control vs TMT, +p<0.05; TMT, n=14, water control, n=6, ethyl vanillin control, n=4). TMT-exposed mice spent less time in the open arms of the EPM on pnd 28 and pnd 41 compared to controls (Fig. 3, C; Control vs TMT, ++p<0.001; TMT, n=14, water control, n=6, ethyl vanillin control, n=4). It is acknowledged that repeat testing on the EPM assesses phobia/fear (35). On pnd 20, pnd 28 and 41, within-group analysis showed no significant differences in the total time spent in the open arms of the EPM across the different test days in the control group. However, within-group analysis of the TMT-exposed group showed a significant difference in time-spent in the open arm of the EPM on pnd 41 compared to pnd 28 (Fig. 3, C; TMT on pnd 28 vs TMT on pnd 41, ###p<0.0001). TMT-exposure during pubertal transition had a long-lasting effect on anxiety and fear/phobia.

**Figure 3.**
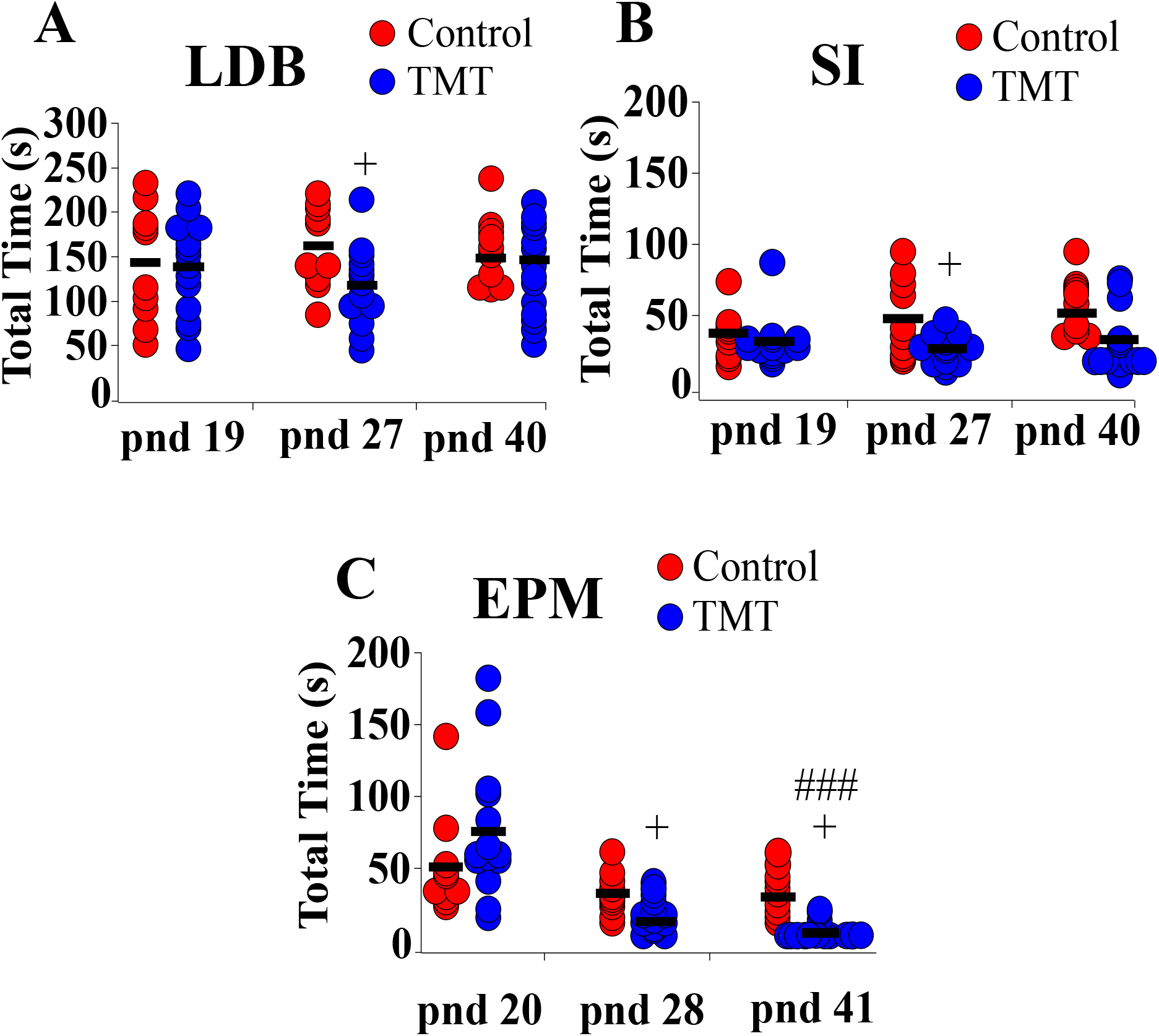
Female C57Bl6/J mice chronically exposed to 2,4,5-Trimethylthiazole (TMT) showed increased anxiety-like behavior on the light-dark box (LDB) and elevated plus maze (EPM) and decreased social interaction with familiar conspecifics on the social interaction (SI) test. (A) Summary of time spent in the light compartment of the LDB on post-natal day (pnd) 19 (before beginning of TMT-exposure), 27 (during TMT exposure) and 40 (after termination of TMT-exposure), (B) time spent socially interacting with familiar conspecifics on pnd 19, 27 and 40, and (C) time spent in the open arm of the EPM on pnd 20, 28 and 41. +p<0.05 time spent on LDB, SI and EPM of control group (combined water control: n=6; ethyl vanillin control: n=4) vs TMT-exposed group (n=14) on pnd 27 or 28 (during TMT-exposure); ###p<0.0001 time spent in the open arm of the EPM in the TMT-exposed group on pnd 28 vs pnd 41.

### Selective expression of DREAD(Gq) in MePD Ucn3 neurons

Evaluation of m-Citrine, hM3D, expression in tdTomato labelled neurons from AAV-injected Ucn3-cre-tdTomato mice revealed that 86 ± 5% of MePD Ucn3 neurons expressed hM3D and the number of tdTomato labelled Ucn3 neurons per side per slice in the MePD was counted at 72.90 ± 5.48 (mean ± SEM) with the number of AAV-hM3D-mCitrine infected neurons being 63.00 ± 6.83 (mean ± SEM) (n=8). A representative example is shown in Fig 4, A-F.

**Figure 4.**
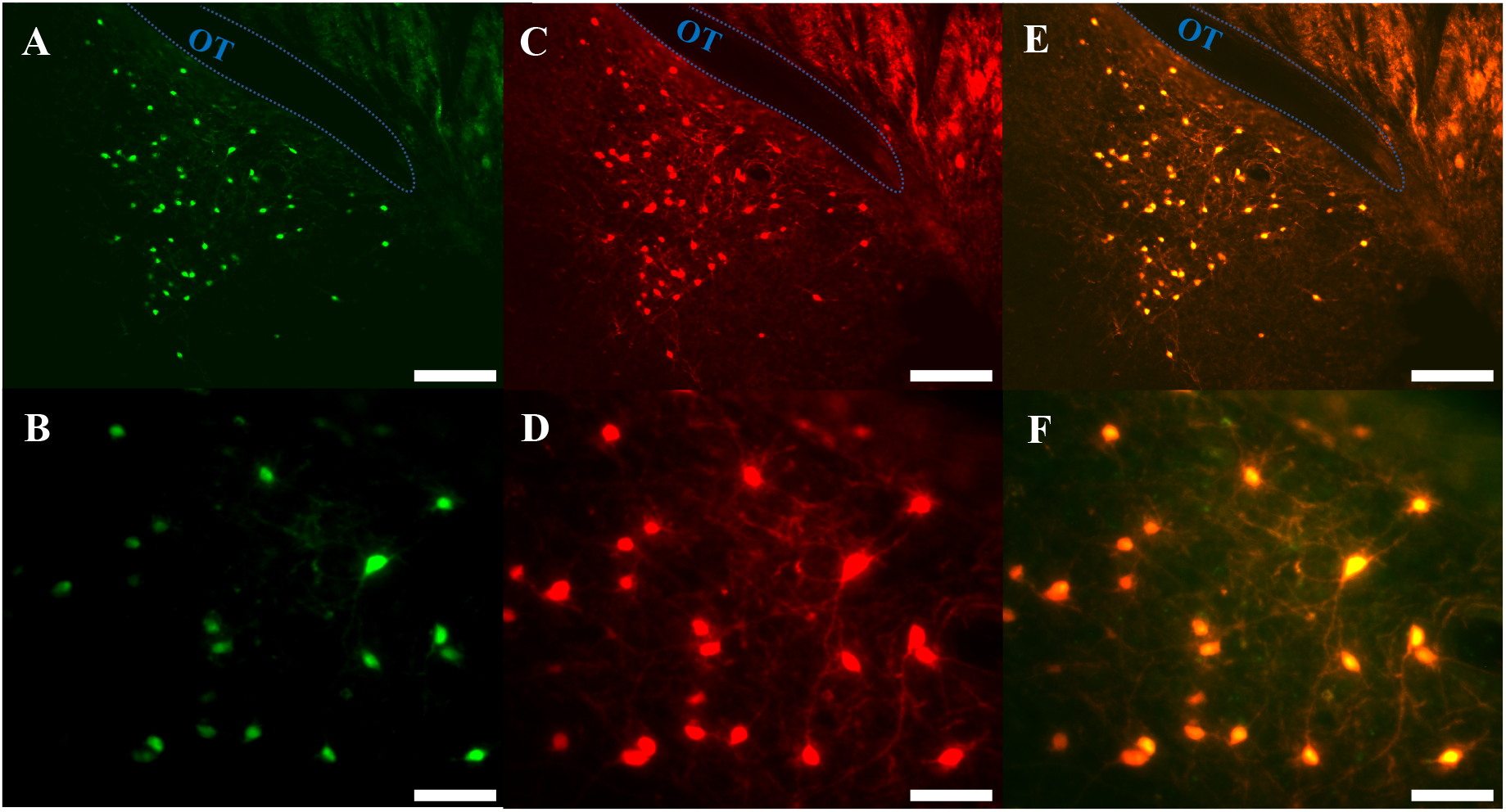
Expression of AAV5-hSyn-DIO-HA-hM3D(Gq)-IRES-mCitrine in posterodorsal medial amygdala (MePD) Urocortin 3 (Ucn3) neurons. (A-F) Representative dual fluorescence photomicrographs of the MePD from a Ucn3-cre-tdTomato female mouse injected with AAV5-hSyn-DIO-HA-hM3D(Gq)-IRES-mCitrine. Ucn3 neurons labelled with m-Citrine (A) and tdTomato (C) appear yellow/orange (E). (B), (D) and (F) are a higher power view of (A), and (E) respectively. Scale bars represent A, C, E 100 µm and B, D, F 25 µm; OT, Optic tract (blue line).

### Selective DREADD activation of Ucn3 neurons in the MePD delays puberty onset

Bilateral DREADD activation of MePD Ucn3 neurons during the pubertal transition period, pnd 21 to 35, of Ucn3-cre-tdTomato mice delayed FE without altering VO compared to non-surgery and AAV-YFP controls (Fig. 5, A and B; Control vs DREADD, +p<0.05; DREADD, n=8, control AAV, n=6, cre-negative control, n=4, no CNO, n=3). Data for cre-negative control, no CNO and control AAV-YFP injected mice were combined as control since there was no significant difference between the control groups. Body weight was unaffected between the experimental groups (Fig. 5, C; Control vs DREADD). These data show that activation of MePD Ucn3 during the pubertal transition delays pubertal timing. Mice with misplaced injections (n=1) were excluded from the analysis.

**Figure 5.**
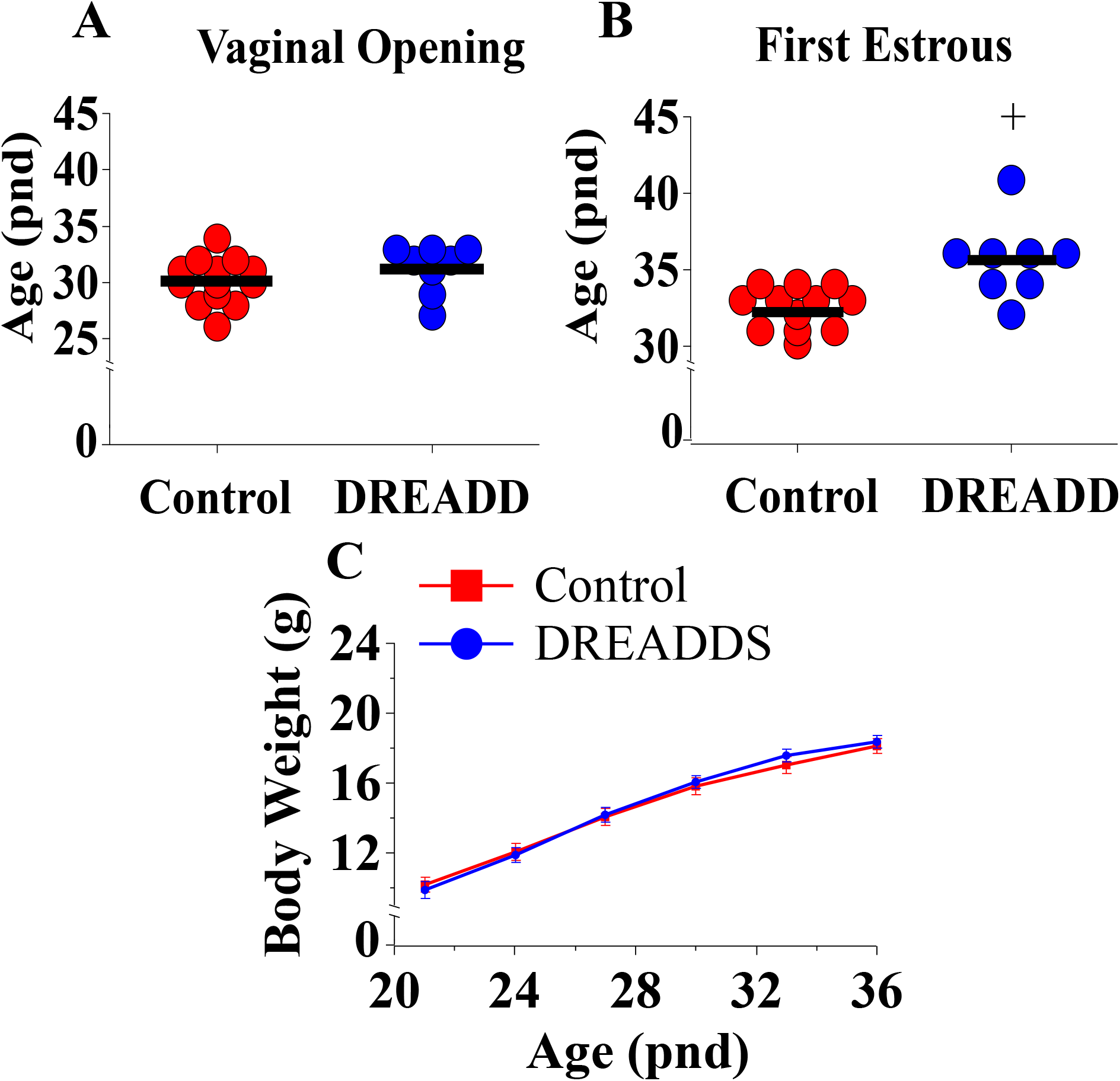
DREADD (hM3D) activation of urocortin 3 (Ucn3) neurons in the posterodorsal sub-nucleus of the medial amygdala (MePD) of Ucn3-cre-tdTomato female mice from post-natal day (pnd) 21 for 14 days delays first estrous (FE) without affecting vaginal opening (VO) and body weight (BW) in urocortin3-cre-tdTomato female mice. Effect of MePD Ucn3 activation on day of VO (A), FE (B) and (C) BW weight gain between pnd 21 and 45. +p<0.05 Control group (combined control AAV: n=6; cre-negative control: n=4; no CNO n=3) vs DREADD group (n=8).

### DREADD activation of Ucn3 neurons in the MePD suppresses pre-pubertal LH pulse frequency

Bilateral DREADD activation of MePD Ucn3 neurons during the pubertal transition period, pnd 21 to 35, of Ucn3-cre-tdTomato mice suppressed pre-pubertal LH pulse frequency sampled on pnd 29 compared to controls (Fig. 6, A, B and C; Control vs DREADD, +p<0.05; DREADD, n=5, control AAV, n=3, cre-negative control, n=3). Data for cre-negative control and control AAV-YFP injected mice were combined as control since there was no significant difference between the control groups. These data show that activation of MePD Ucn3 during pubertal transition suppresses pre-pubertal pulsatile LH secretion. Mice with misplaced injections (n=1) were excluded from the analysis.

**Figure 6.**
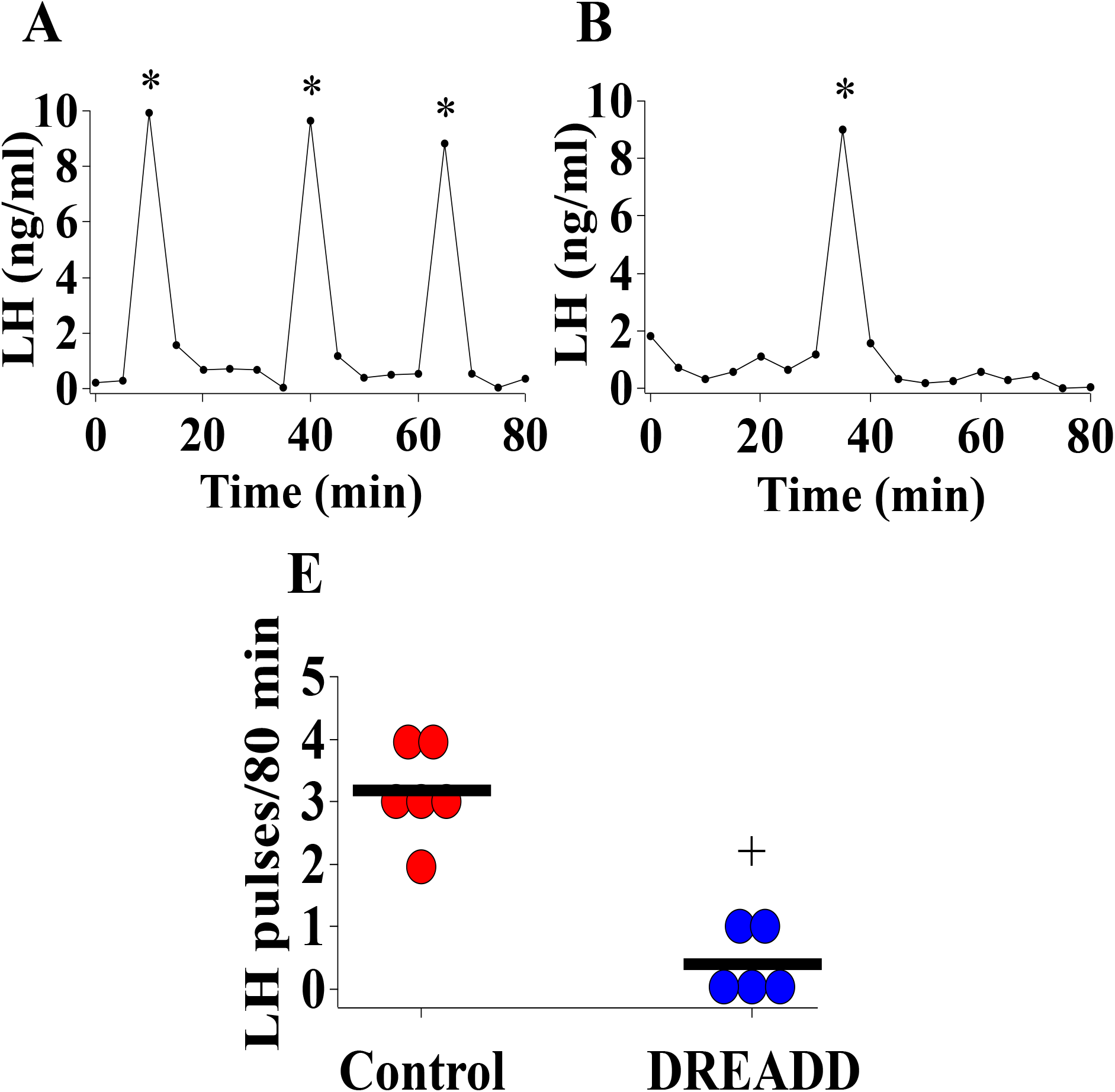
DREADD (hM3D) activation of urocortin 3 (Ucn3) neurons in the posterodorsal sub-nucleus of the medial amygdala (MePD) in Ucn3-cre-tdTomato female mice from post-natal day (pnd) 21 for 14 days suppresses pre-pubertal LH pulse frequency sampled on pnd 29. Representative LH pulse profile from Ucn3-cre-tdTomato female mice on pnd 29 with (A) control mouse, (B) DREADD (hM3D) activated mouse, (C) Summary of LH pulse frequency on pnd 29 of control and DREADD group during the 80 min blood sampling period. LH pulses detected by the DynePeak algorithm are indicated with an asterisk located above each pulse on the representative LH pulse profiles. +p<0.05 Control group (combined control AAV: n=3; cre-negative control: n=3) vs DREADD group (n=5).

## Discussion

The present study shows for the first time that chronic exposure to the psychogenic stressor, predator odor, from pnd 21 for 14 days suppresses the pubertal acceleration of LH pulse frequency, delaying puberty onset, while concurrently inducing long-lasting anxiety and fear-like behaviour. Moreover, DREADD activation of MePD Ucn3 neurons during this same developmental period delays puberty onset while suppressing pre-pubertal LH pulse frequency, demonstrating that MePD Ucn3 signalling may play an important role in stress-related changes in pubertal timing.

Hypothalamic kiss1 signalling is pivotal in regulating puberty onset in various species, including rodents and humans (7,11). Pubertal onset is thought to be timed by an increase in GnRH pulse generator frequency, however the mechanisms underlying the timing of ARC kiss1 neuronal activation to trigger pubertal onset are not well established. Nevertheless, puberty onset has been associated with increased ARC Tac2 (encoding NKB) and kiss1 mRNA expression in mice (36,37), which are critical components of the GnRH pulse generator and there is an increase in ARC kiss1 promoter activity during pubertal development (38). The amygdala is known to influence pubertal timing in primates (17), and more specifically, the MePD has been shown to regulate pubertal timing and reproductive function in rodents (4,19), vis-à-vis MePD kiss1 receptor antagonism delays puberty, reduces the occurrence of preovulatory LH surges in rats (25) and MePD kiss1 neuronal activation heightens sexual partner preference in male mice (39). Recently, we have shown optogenetic stimulation of MePD kiss1 neurons increases GnRH pulse generator frequency in adult female mice (26) and preliminary data from our lab shows that activation of MePD kiss1 neurons during pubertal transition advances the onset of first estrous, thus MePD kiss1 neurons may be an upstream regulator of pubertal timing (40).

We explored the effect of chronic psychological stress exposure on pubertal timing, pre-pubertal GnRH pulse generator activity and anxiety-like behaviour. Stress exposure during the juvenile period is known to alter pubertal timing and induce anxiety behaviour in mammals, including rodents, primates and humans (1,2,4,41). Predator odor exposure in rodents is a model for PTSD; increasing fear and anxiety behaviour as well as reducing social interaction (3). Previously, we have shown that acute TMT exposure suppresses LH pulsatility (5) and restraint stress, another psychological stressor, inhibits LH pulse frequency while reducing ARC kiss1 cfos expression in adult mice (42). Moreover, we have shown that cat odor delays puberty and induces a fear response in rats (4). In the present study we show that the predator odor, TMT, reduces LH pulsatility and delays the first estrous in female mice. In mice, VO and FE do not coincide, wherein VO is not tightly coupled to the first ovulation, but rather a variable time gap exists between VO and FE; the latter associated with first ovulation and considered the unequivocal marker of puberty in this species (43). Additionally, pre-pubertal LH pulse frequency in the control group tended to increase potentially marking the acceleration of the GnRH pulse generator as the time of puberty onset approaches. Contrastingly, pre-pubertal LH pulse frequency in the TMT-exposed group was significantly lower compared to controls and we did not observe an increase in GnRH pulse generator activity between pnd 26 and 29.

Puberty is a sensitive period where enhanced neuroplasticity provides a context for the impact of stress on the development of long-term anxiety disorders in humans (44). Early-life stress increases anxiety-like behaviour in juvenile and adult mice linked to hyper-connectivity between fronto-limbic circuits, involving the amygdala (45). The medial amygdala is activated in response to external stressors and electrophysiological recordings reveal that predator urine robustly activates the MePD, which is associated with delaying pubertal onset (4) and supressing LH pulsatility in rodents (5,46). We found that predator odor-induced pubertal delay was accompanied by increased anxiety-like behaviour, with mice spending significantly less time in the light compartment of the LDB and in the open arms of the EPM during stress exposure. Moreover, we observed that mice exposed to TMT exhibited long lasting fear-like behaviour where they spent significantly less time in the open arm of the EPM compared to controls on pnd 41; 6 days after termination of stress exposure. This is consistent with our previous studies showing rats exposed to cat odor during pubertal development display increased long-lasting fear-like behaviour on the EPM even after the termination of stress exposure (4). Moreover, exposure to fox odor induces long-lasting anxiety on the EPM in rats (47), while mice exposed to foot-shock stress during adolescence exhibit increased startle reflexes into adulthood indicative of lasting anxiety-like behaviour (48). Additionally, TMT-exposed mice showed significantly reduced social interaction with familiar con-specifics during stress exposure whereas sociability tended to increase along with reproductive maturation in the control group, which is consistent with our previous observations where exposure to cat odor during puberty decreased sociability in rats (4).

Central administration of CRF inhibits kiss1 expression in the ARC and POA in rats (49). In women with functional hypothalamic amenorrhea there is a correlation between decreased serum kisspeptin and increased CRF concentrations (50). Moreover, an endogenous CRF tone regulates pubertal timing where central administration of CRF in pnd 28 rats delays puberty onset and administration of astressin-B, a CRFR1 antagonist, advances puberty (28). Central antagonism of CRFR2 blocks the suppressive effect of stress on LH pulsatility in rodents (51). These data indicate stress neuropeptides may provide an inhibitory tone on the timing of puberty. Predator odor activates CRFR2 positive neurons in the medial amygdala of rats (52) and we have recently shown that TMT-induced suppression of LH pulsatility is mediated by MePD Ucn3 and CRFR2 signalling in mice (5). In the present study, selective activation of MePD Ucn3 neurons during pubertal development delayed puberty and suppressed pre-pubertal LH pulse frequency, demonstrating that MePD Ucn3 signalling plays a key role in regulating pubertal timing.

The MePD has a major GABAergic output to key hypothalamic reproductive nuclei (21,53) and we know the amygdala exerts an inhibitory brake on pubertal timing (17,19). Ucn3 neurons in the medial amygdala have an interneuron-like appearance (54) where Ucn3 fibres overlap with CRFR2 expression (55,56) and we have shown that antagonism of MePD CRFR2 blocks the suppressive effect of TMT on LH pulse frequency in adult mice (5), thus Ucn3 neurons possibly connect to and signal via CRFR2 within the MePD. Moreover, the majority of MePD CRFR2 neurons co-express GAD65 and 67 (29) indicating they are GABAergic and may be involved in mediating stress-induced suppression of the GnRH pulse generator (22,53). Therefore, MePD Ucn3 neurone activation may modulate pubertal timing possibly via enhancing GABAergic output to inhibit GnRH pulse generator activity (5,21,23).

Our findings show for the first time that early life exposure to a psychosocial stressor disrupts GnRH pulse generator frequency to delay puberty, and Ucn3 signalling in the MePD may be involved in mediating this response. These findings provide novel insight into the key interactions between the emotional stress centres and reproductive centres in the brain that control pubertal onset in response to the external environment. Alterations in the timed onset of puberty has major implications later in life with increased risk of diverse adverse outcomes, including cancer, gynaecologic/obstetric, cardio-metabolic and neuro-cognitive categories in humans (57). Understanding the mechanisms involved in controlling pubertal timing and the impact of stress is crucial to ultimately aid future development of more effective treatments for stress-related disorders of puberty.

## Acknowledgements

The authors gratefully acknowledge the financial support from UKRI: BBSRC (BB/S000550/1) and MRC (MR/N022637/1). DI is a PhD student funded by MRC-DTP studentship at King’s College London.

## Data Availability Statement

The original contributions presented in the study are included in the article/supplementary material. Further inquiries can be directed to the corresponding author.

## Ethics Statement

The animal study was reviewed and approved by the Animal Welfare and Ethical Review Body Committee at King’s College London.

